# Localization of muscarinic acetylcholine receptor dependent rhythm generating modules in the *Drosophila* larval locomotor network

**DOI:** 10.1101/2021.03.08.432150

**Authors:** Julius Jonaitis, James MacLeod, Stefan R. Pulver

## Abstract

Mechanisms of rhythm generation have been extensively studied in motor systems that control locomotion over terrain in limbed animals; however, much less is known about rhythm generation in soft-bodied terrestrial animals. Here we explored how muscarinic acetylcholine receptor (mAChR) dependent rhythm generating networks are distributed in the central nervous system (CNS) of soft-bodied *Drosophila* larvae. We measured fictive motor patterns in isolated CNS preparations using a combination of Ca^2+^ imaging and electrophysiology while manipulating mAChR signalling pharmacologically. Bath application of the mAChR agonist oxotremorine potentiated rhythm generation in distal regions of the isolated CNS, whereas application of the mAChR antagonist scopolamine suppressed rhythm generation in these regions. Oxotremorine raised baseline Ca^2+^ levels and potentiated rhythmic activity in isolated posterior abdominal CNS segments as well as isolated anterior brain and thoracic regions, but did not induce rhythmic activity in isolated anterior abdominal segments. Bath application of scopolamine to reduced preparations lowered baseline Ca^2+^ levels and abolished rhythmic activity. These results suggest the presence of a bimodal gradient of rhythmogenicity in the larval CNS, with mAChR dependent rhythm generating networks in distal regions separated by medial segments with severely reduced rhythmogenic abilities. This work furthers our understanding of motor control in soft-bodied locomotion and provides a foundation for study of rhythm generating networks in an emerging genetically tractable locomotor system.

## Introduction

Understanding how and where rhythms are generated in motor systems is fundamental to understanding how animals move. The operation and location of central pattern generating (CPG) networks controlling terrestrial locomotion has been extensively studied in limbed animals such as mice (Meehan et al., 2012; Hägglund et al., 2013), cats (Andersson and Grillner, 1981; Hultborn and Nielsen, 2007), turtles (Guertin and Hounsgaard, 1998), crustaceans (Mulloney and Smarandache-Wellmann, 2012; Zhang et al., 2014) and insects (Blaesing and Cruse, 2004; Mantziaris et al., 2017, 2020). In contrast, much less is known about CPG networks in soft-bodied terrestrial animals. Soft bodied crawlers are intriguing from a motor control perspective; their fluid and air filled semi-compressible bodies have high degrees of freedom of movement and present a unique set of motor control challenges that are different from those faced by limbed animals (van Griethuijsen and Trimmer, 2010, 2014; Heckscher et al., 2012).

Work in leeches, caterpillars and nematodes has provided important initial insights into the organization of CPG networks controlling soft-bodied crawling. In both leeches and caterpillars, there is strong evidence that crawling CPGs are organized as chains of coupled segmental oscillators (Johnston and Levine, 1996; Puhl and Mesce, 2008). In contrast, nematode crawling CPG circuits appear to be more continuous with circuits spread across segmental boundaries (Fouad et al., 2018; Gao et al., 2018; Wen et al., 2018). In all three systems, neuromodulators play critical roles in reconfiguring locomotor circuits to select crawling over other motor programmes (Johnston and Levine, 1996; Puhl and Mesce, 2008; Vidal-Gadea et al., 2011).

The *Drosophila* larval locomotor system presents a unique opportunity to further study the neural control of locomotion in a genetically tractable soft bodied animal. Recent work in this system has focused on characterizing the functional roles of identified interneurons, and on uncovering functional circuit motifs based on EM reconstructions and selective activation and inhibition of interneuron subtypes in the CNS (Kohsaka et al., 2014, 2019; Fushiki et al., 2016; Takagi et al., 2017; Carreira-Rosario et al., 2018; Tastekin et al., 2018; Heckscher et al., 2015). Previous work has revealed that the isolated larval CNS generates fictive motor programmes for forward locomotion, backward locomotion and head sweeps (Pulver et al., 2015) and provided initial insights into the location of CPG networks (Berni et al., 2012; Berni, 2015; Sims et al., 2019). Computational studies have modelled the larval VNC as comprised of oscillators in each hemi-segment (Gjorgjieva et al., 2013) or as recurrent networks (Zarin et al., 2019); however, the precise location and neuromodulatory requirements for rhythm generation in this system have remained unclear.

Signalling through metabotropic muscarinic receptors (mAChRs) is critical for rhythm generation in a variety of insect locomotor networks (Ryckebusch and Laurent, 1993; Büschges et al., 1995; Johnston and Levine, 2002; Buhl et al., 2008). The underlying genes and pharmacological profiles of 3 separate *Drosophila* mAChR variants have been well characterized, with two variants (A and B type) showing similar pharmacological profiles to orthologous receptors in vertebrates (Collin et al., 2013; Ren et al., 2015). Recent work has also characterized the behavioural and physiological effects of inhibiting expression of mAChRs in defined neuronal populations using RNA interference (Malloy et al., 2019). Although powerful, these genetic manipulations operate over the time scale of days, leaving open the possibility of compensatory changes over developmental time. To date no studies have directly explored the role(s) that acute manipulation of mAChR signalling plays in modulating CPG activity in defined regions of the larval locomotor system.

Here we use a combination of physiology, microsurgery and pharmacology to explore the basic organization of muscarinic dependent rhythm generation in the larval VNC. We show how mAChR dependent rhythm generating modules are distributed across the *Drosophila* larval VNC and provide evidence for the presence of multiple rhythm generating regions, as well as gradients of rhythmogenicity across the anterior-posterior axis of the larval CNS. This work provides a foundation for unravelling the cellular mechanisms of rhythm generation in the *Drosophila* larval locomotor system.

## Methods and Materials

### Animal rearing and genetic constructs

Flies were grown on standard corn-meal based fly food. Animals were maintained in temperature controlled incubators at 21 - 22° C with ∼55-60% humidity. The GAL4-UAS system (Brand and Perrimon 1993) was used to drive expression of the Ca^2+^ indicator GCaMP6s(Chen et al., 2013) in all neurons. We used 57C10-GAL4 inserted in the attP2 landing site for all experiments. This GAL4 incorporates the promoter fragment for a common presynaptic protein (synaptobrevin)) and has been used in previous pan-neuronal imaging studies (Lemon et al., 2015; Pulver et al., 2015). We used a construct that has 57C10-GAL4 combined with UAS-GCAMP6s inserted in the attP40 landing site.

### Dissection

Feeding 3^rd^ instar larvae were used for all experiments. In each experiment, single larvae were placed in an acrylic recording chamber lined with a thin layer (1 mm) of SYLGARD 184 Silicone Elastomere (Sigma-Aldrich, Irvine, United Kingdom). Each larva was placed in the centre of the chamber, dorsal side up and pinned to the substrate by placing one insect pin in the mouth hook region and one pin in the most posterior abdominal segments (Fig. 1A). A dorsal incision was then made the length of the body using microdissection scissors (Fig. 1B). Four fine tungsten wire pins (0.025 mm, Alfa Aesar, Fisher Scientific) were then used to pin the larval body wall flat onto the surface of the dish and internal organs of the larvae were then carefully removed, leaving the CNS attached to the body wall (Fig. 1C). The CNS including the brain lobes and the ventral nerve cord (VNC), was then carefully cut out and placed away from the larval body carcass. The isolated CNS was pinned out dorsal side up by placing 2-4 fine tungsten wire pins around the preparation (California Fine Wire, Grover Beach, CA). Care was taken not to damage the brain lobes or VNC. In some experiments, different regions of the CNS were ablated (Fig. 1D). Ablations were made using micro dissecting scissors and/or sharpened hypodermic needles (Sigma-Aldrich, United Kingdom). Before proceeding with recordings from ablated preparations, remaining nerve roots were counted to determine which segments were still intact and which had been removed. Preparations that showed clear evidence of structural damage away from the immediate region where the cut was made were excluded from analysis. During all dissections animals were submerged in physiological saline containing (in mM) 135 NaCl, 5 KCl, 2 CaCl^2^, 4 MgCl^2^, 5 TES, and 36 sucrose, pH 7.15. During dissection the saline was exchanged 2-3 times to remove debris.

**Figure 1.**
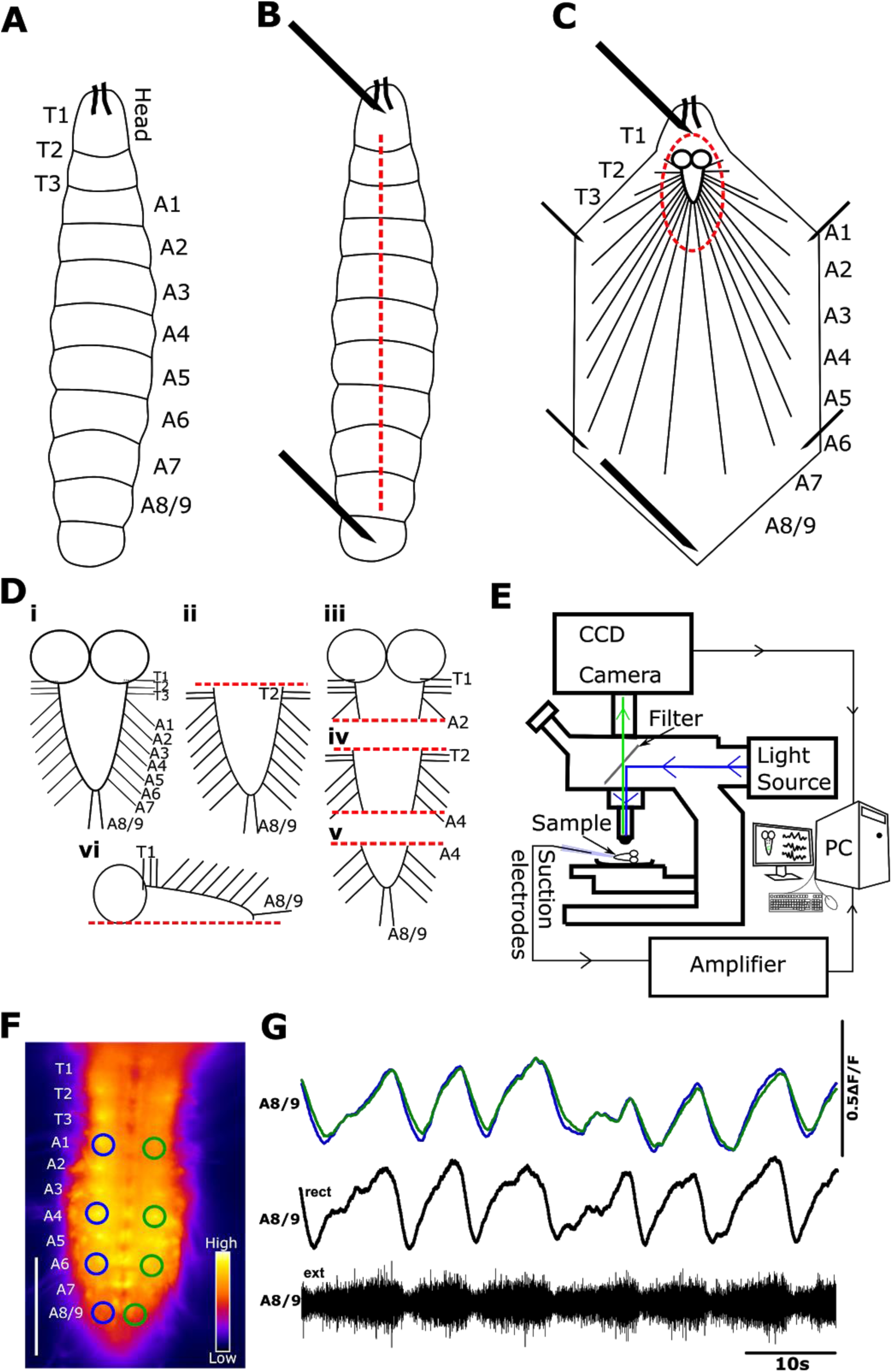
*Drosophila* larval preparation for recording fictive locomotor patterns. **A**) Schematic of the body of a 3^rd^ instar *Drosophila* larva. T1-T3 denote larval thoracic segments, A1-A8/A9 denote abdominal segments. **B**) Schematic showing first steps of dissection. Head and tail of the larva are secured with insect pins. Dashed red line indicates where dorsal incision is made with micro-dissection scissors. **C**) Schematic of larval body filleted with CNS exposed and ready for removal. Dashed lines indicate where cuts are made to remove the CNS. **D**) Schematic of isolated CNS and cuts made to isolate different CNS regions. i) Intact CNS ii) Brain lobes removed, iii) abdominal segments removed, iv) Brain lobes and distal regions removed, v) brain lobes and anterior regions removed vi) One side of CNS removed. **E**) Experimental setup for simultaneous Ca^2+^ imaging and electrophysiological recordings. **F)** Single frame showing raw fluorescence from an experiment in which Ca^2+^ activity is imaged panneuronally. ROI size and placement is shown on the right (green) and left (blue) hemi-segments, along with segment designations (to left of image). Differences in fluorescence intensity are indicated by color lookup table bar at bottom right. Scale bar, 100μm. **G)** Percent change in fluorescence within A8/9 ROIs shown in (F) (scale bar shows 50% ΔF/F). Simultaneous extracellular nerve root recording from A8/9 confirms that Ca^2+^ signals from ROIs reflect motor output from hemi-segments.

### Electrophysiology

In a subset of experiments, extracellular suction electrode recordings from motor nerve roots projecting from the VNC were performed in order to record fictive motor patterns. Borosilicate glass capillary tubes (Kwik-Fil 1B150F-4, World Precision Instruments, Hitchin, United Kingdom) were pulled using a Model P90 micropipette puller (Sutter instruments, Hitchin, United Kingdom) and the tips were broken to match the diameter of a single nerve root. Three electrodes were then maneuvered towards the isolated CNS using a MP-285 (Sutter Instruments) mechanical micromanipulator and two manual manipulators (DT3-100 XYZ and DT3-130 XYZ, Siskiyou, Grants Pass, Oregon). Single nerve roots of interest were then drawn into the suction electrodes. Typically, one electrode was placed on a posterior nerve root (A7 or A8/9) and two placed contralateral anterior nerve roots (T3 or A1) Electrophysiological signals were amplified using a Model 1700 extracellular amplifier (A-M Systems, Sequim, WA), digitized using a model NI-USB-6229 data acquisition board (National Instruments, London, United Kingdom) and recorded using Winfluor V4.0.3 (University of Strathclyde, Glasgow, United Kingdom). Recordings were then analyzed offline using Spike2 Version 7.08b (Cambridge Electronic Design Limited, Milton, United Kingdom) or Dataview (Bill Heitler, University of St Andrews, United Kingdom).

### Ca^2+^ imaging in the isolated CNS

Isolated CNS preparations expressing GCaMP6s panneuronally were imaged using a modified BX51 epifluorescence microscope (Olympus, Japan). An OptoLED (Cairn Instruments, Kent, UK) was used to generate blue light (470 nm), which was directed to the preparation using a dichroic mirror in a custom built housing (Cairn Instruments, Kent, UK). Emitted light was filtered using a green fluorescent protein (GFP) emission filter. Images were captured with an iXon3 EMCCD camera (Andor Instruments, Belfast, United Kingdom)(Fig. 1E). Preparations were recorded for 25 - 30 minutes (depending on the experimental protocol) using a N20X-PFH 20x water immersion objective (Olympus, Japan). Images were typically acquired at 9.94 frames per second with a 100 ms exposure time using WinFluor. Changes in fluorescence values were extracted from regions of interest (ROIs) in thoracic (T2-3) and abdominal (A2, 4 and 6) segments (Fig. 1F) using Fiji or ImageJ (Schindelin et al., 2012; Schneider et al., 2012) available online (https://imagej.net/). Extracted ROI fluorescence signals were visualized as percentage change in fluorescence from average values (ΔF/F) using custom scripts written in Python (Python Software Foundation, www.python.org).

### Application of pharmacological agents

A custom built superfusion system incorporating a U120 peristaltic pump (Watson-Marlow Pumps, Falmouth, United Kingdom) was used to superfuse physiological saline over preparations at a rate of 0.1 ml/s. Drugs were bath applied and washed out by switching between different gravity fed inflows. Control periods typically lasted 5 - 7 minutes, drug applications lasted 6 minutes, and wash periods lasted 6 - 10 minutes. It typically took ∼45 - 60 seconds for full bath exchange after switching feeds. In ablation experiments, we typically waited at least 8 - 10 minutes after each cut before conducting experiments to reduce the chances of artifacts due to injury firing. The muscarinic agonist oxotremorine (Sigma-Aldrich, Irvine, United Kingdom) was applied at concentrations ranging from 10^-4^ to 10^-6^M. The muscarinic antagonist scopolamine (Sigma-Aldrich, Irvine, United Kingdom) was applied at concentrations ranging from 3 x 10^-4^ to 3 x 10^-7^M.

### Waveform analysis, statistics and figure making

Event detection and waveform analysis was carried out using custom scripts in Spike 2 and Dataview. Statistical tests were carried out in GraphPad Prism 5 (GraphPad, San Diego, California). 1-way repeated measures ANOVA with Bonferroni multiple comparison post-hoc tests were used to make statistical comparisons. When datasets failed the Shapiro-Wilk normality test (p < 0.05), non-parametric Friedman with Dunn’s post hoc multiple comparison tests were used. Figures were made in GraphPad Prism 5 or 8 and finalized in Inkscape (available free online at https://inkscape.org/)

## Results

To establish the fundamental features of larval locomotor rhythms, we recorded fictive locomotor activity using Ca^2+^ imaging combined with electrophysiological recordings from nerve roots in 3rd instar larval VNCs (Fig. 1G and 2A, B). Imaging Ca^2+^ panneuronally using a highly sensitive GCaMP variant (6s) while also recording from multiple nerve roots maximized chances of detecting rhythmic activity in both interneuron and motor neuron populations. Reading out Ca^2+^ signals in each hemi-segment of the VNC provided a summed measure of all activity within that hemi-segment; overlaying signals corresponding segments across the midline allowed visualization of bilaterally symmetric and asymmetric events (Fig. 2A-C and Fig. 3A, B green and blue traces). Subtracting signals from corresponding ROIs across the midline allowed visualization of bilaterally asymmetric activity patterns (Fig. 2A, 3A gray trace at top). Simultaneous extracellular recordings were performed from multiple nerve roots to confirm that Ca^2+^ signals in the panneuronal expression pattern in the larval VNC corresponded to motor activity (Fig. 1F, G and 2B, C).

**Figure 2.**
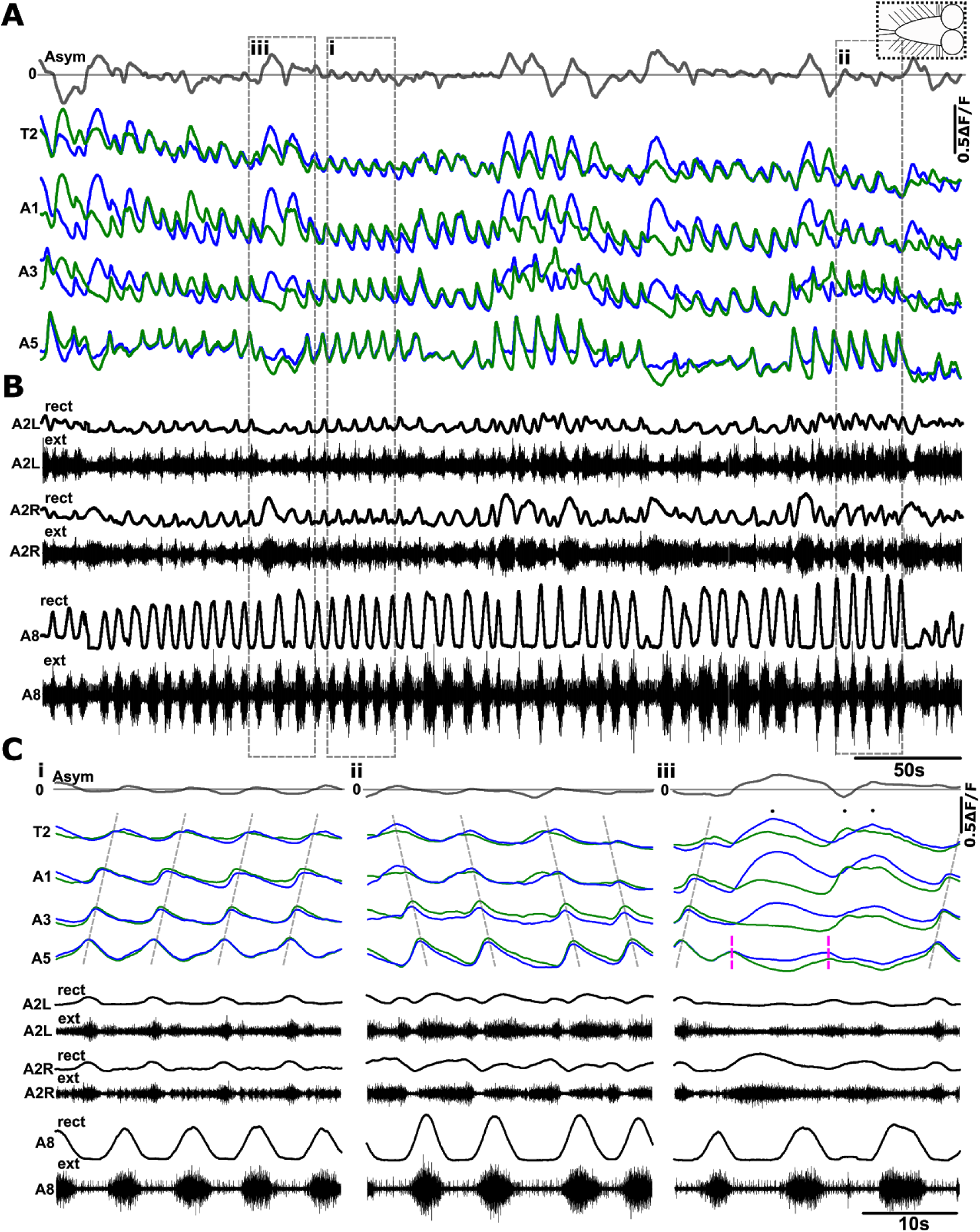
Spontaneous activity in the isolated larval CNS. **A**) Percent change in fluorescence (ΔF/F) extracted from ROIs placed on different hemi-segments on left (blue) and right (green) sides of the VNC. Bilateral asymmetry (Asym) in Ca^2+^ intensity (top dark grey trace) indicated between T2 left and right sides. Peaks above 0 baseline (light grey line) indicate higher Ca^2+^ intensity on the left side and below zero indicate higher Ca^2+^ intensity on the right side of the VNC. Vertical scale bar indicates 50% ΔF/F. Dashed box at the top right corner shows schematic of type of CNS preparation used in experiment. **B**) Simultaneous extracellular (ext) suction electrode recordings from three nerve roots (black traces), abdominal segment left and right sides (A2L and A2R) and single nerve root recording from abdominal segment 8 (A8). Extracellular nerve signals recorded from each electrode were rectified and smoothed by filtering with a moving average filter with a time constant of 0.9 s (rect). C**i**) Expanded view of both Ca^2+^ signals and suction electrode recordings showing forward waves (dashed grey line). C**ii**) examples of backward waves. C**iii**) examples of bilateral asymmetries (black dots over events). Posterior bursts that did not propagate into forward waves are shown with magenta dashed lines.

**Figure 3.**
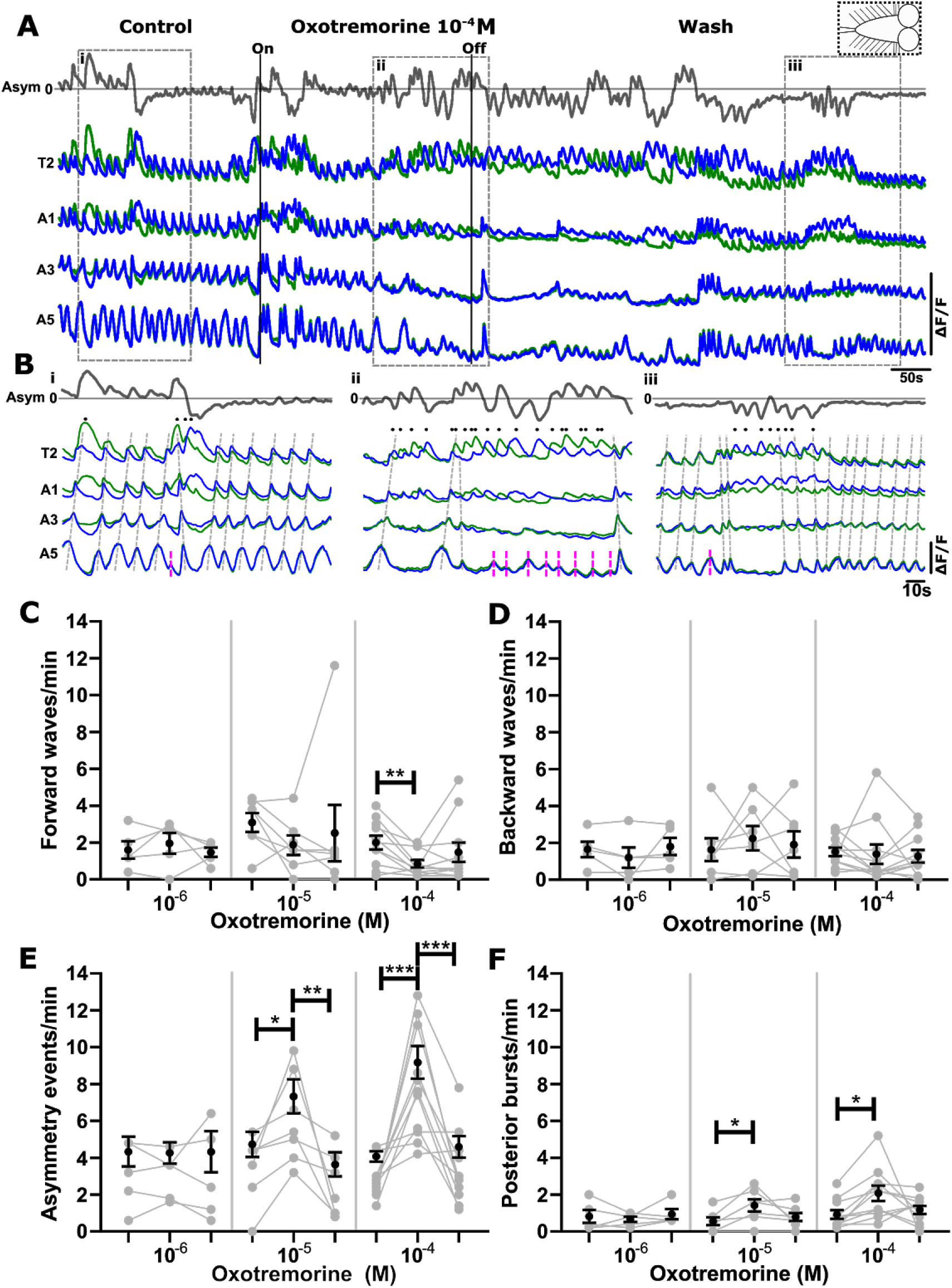
The muscarinic agonist oxotremorine increases frequency of bilaterally asymmetric activity in anterior regions and bilaterally symmetric bursts in posterior regions. **A**) Percent change in fluorescence (ΔF/F) extracted from ROIs placed on different segments on left (blue) and right (green) sides of the VNC. Bilateral asymmetry (Asym) in Ca^2+^ intensity (top dark grey trace) indicated between T2 left and right sides. Dashed box at the top right corner shows schematic of type of CNS preparation used in experiment. **Bi**) Expanded view of Ca^2+^ signals during control period, B**ii**) expanded view of Ca^2+^ signals during oxotremorine 10^-4^M application, B**iii**) expanded view of Ca^2+^ signals in wash period. **C-F**) Mean +/− S.E.M events per minute of different fictive motor patterns produced before (light grey bars), during (black bars) and after (dark grey bars) application of different concentrations (M) of oxotremorine. Asterisks indicate significant differences between different conditions of the experiment (one-way repeated measures ANOVA with Bonferroni multiple comparison post-hoc test; when datasets failed the Shapiro-Wilk normality test (p < 0.05), non-parametric Friedman with Dunn’s post hoc multiple comparison was used (**C, D:** 10^-4^M, 10^-6^M, **E:** 10^-6^M, **F:** 10^-5^M). *p < 0.05, **p < 0.005, ***p < 0.0001. Sample sizes: n = 5 for 10^-6^M, n = 7 for 10^-5^M and n = 11 for 10^-4^M).

The isolated CNS produced four distinct rhythmic fictive motor patterns in physiological saline (Fig. 2A - C), which mirror patterns of actual activity seen in intact animals(Pulver et al., 2015). The four patterns were: 1) fictive forward waves (defined as bilaterally symmetric Ca^2+^ activity that started in posterior most segments of the VNC and propagated to anterior most segments (Fig. 2Ci); 2) fictive backward waves (defined as bilaterally symmetric Ca^2+^ activity that started in anterior most segments and propagated to posterior most segments (Fig. 2Cii); 3) bilaterally asymmetric activity, i.e. fictive head sweeps (defined as synchronous Ca^2+^ activity peaks in anterior segments that occurred only on one side of the VNC (Fig. 2Ciii, indicated with black dots); 4) posterior bursts, (bilaterally symmetric bursts of activity in distal regions that do not generate waves). These events were termed ‘aborted waves’ in a previous publication because they show qualitative similarities to locomotor waves that initiate but fail to propagate through medial segments (Pulver et al., 2015) (Fig. 2, Ciii, magenta dashed lines).

As a first step, to assess whether muscarinic signalling modulates rhythmic activity in the larval CNS, we bath-applied the muscarinic acetylcholine receptor (mAChR) agonist oxotremorine and measured resulting changes in fictive motor patterns in the isolated CNS (Fig. 3A, B). We used concentrations (10^-6^ - 10^-4^M) that have been shown to activate both A type and B type *Drosophila* mAChRs in previous work (Cattaert and Birman, 2001; Collin et al., 2013; Ren et al., 2015). Bath applied oxotremorine increased the frequency of bilaterally asymmetric events in anterior regions of the VNC in a dose dependent manner (P < 0.0001 in 10^-4^M; P < 0.05 in 10^-5^M; Fig. 3B, E). The two highest concentrations (10^-5^ and 10^-4^M) also significantly increased the frequency of bilaterally symmetric bursts in posterior regions (P < 0.05 in both 10^-4^ and 10^-5^ M, Fig. 3B, F). Forward wave frequencies were also reduced significantly at the highest concentration of oxotremorine (P < 0.005) (Fig. 3C, 10^-4^M). Backward wave frequencies were not significantly affected at any concentrations of oxotremorine Fig. 3B, D). In all experiments, drug application was largely reversible, with most preparations returning towards activity levels seen in control conditions (note grey scatter plots in Fig. 3).

To test if muscarinic signalling is required for rhythm generation, we bath-applied the non-selective mAChR antagonist scopolamine to the isolated intact CNS and measured resulting changes in activity patterns (Fig. 4A, B). We observed significant decreases in forward waves, backward waves, and fictive head sweep frequencies in 3 x 10^-4^ M scopolamine (p < 0.05, p < 0.05, p < 0.001, respectively, Fig. 4C - E) with little to no recovery in the wash periods. There was no significant decrease in forward and backward wave frequencies at lower oxotremorine concentrations; however, scopolamine decreased the frequency of bilateral asymmetries at 3 x 10^-6^M (p < 0.005) and showed a similar albeit, non-significant trend at 3 x 10^-5^M (p > 0.05). Scopolamine had no effect on posterior burst frequency at any concentration (p > 0.05) (Fig. 4C, D, F).

**Figure 4.**
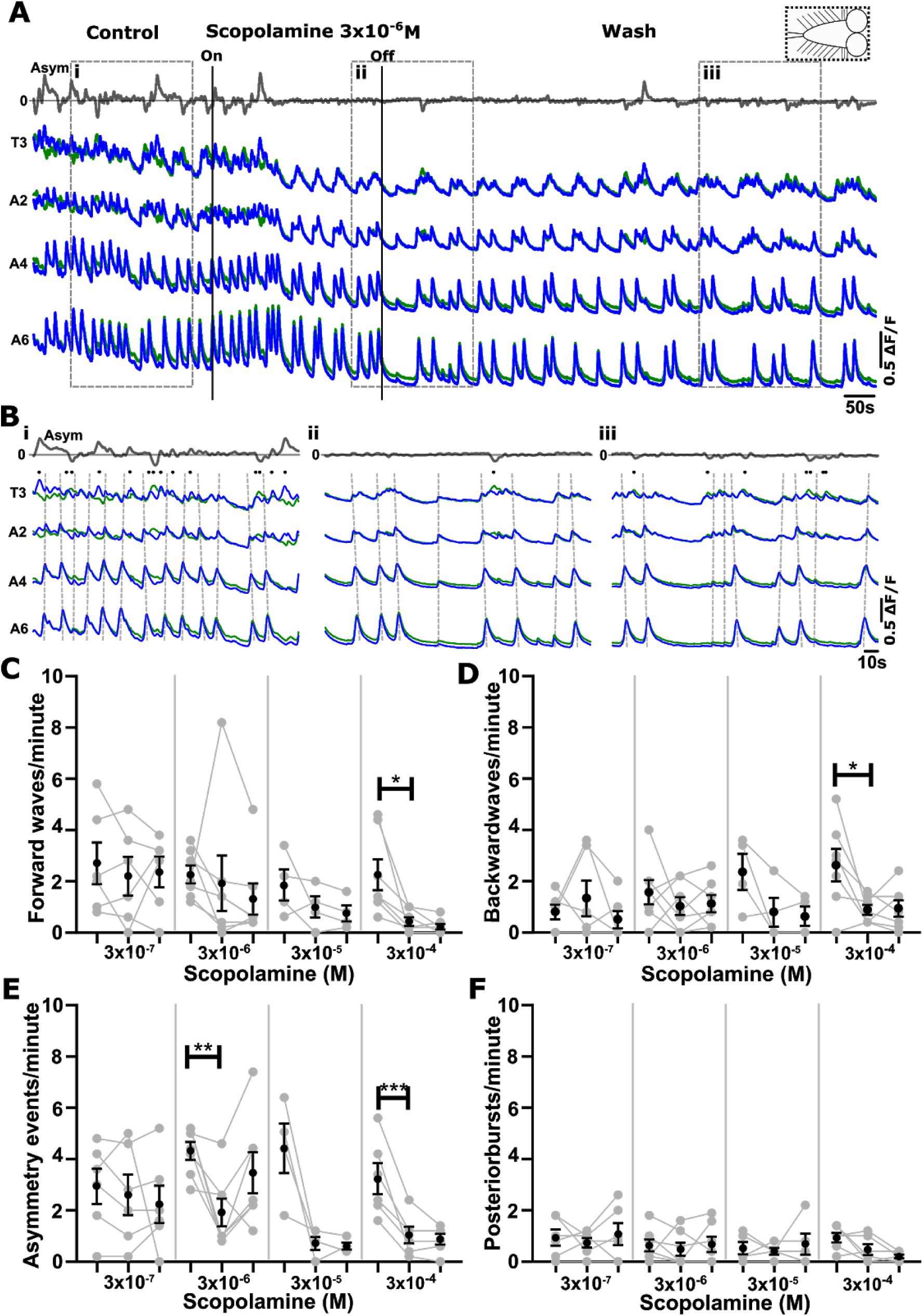
The muscarinic antagonist scopolamine inhibits fictive motor patterns in the isolated CNS. **A**) Ca^2+^ signals in an isolated CNS preparation during application of 3 x 10^-6^M scopolamine. Gray trace at top represents subtraction of left and right side signals in T3. Dashed box at the top right corner shows schematic of CNS sections present in experiment. **Bi**) Expanded view of Ca^2+^ signals during control period, **Bii**) expanded view of Ca^2+^ signals during scopolamine application, **Biii**) expanded view of Ca^2+^ signals in wash period. **C - F**) Mean +/− S.E.M events per minute of different fictive motor patterns produced before, during and after application of different concentrations of scopolamine. Asterisks indicate significant differences amongst groups (*p < 0.05, **p < 0.005, ***p < 0.0001; one-way repeated measures ANOVA with Bonferroni post-hoc test. For datasets that failed the Shapiro-Wilk normality test, non-parametric Friedman tests with Dunn’s post hoc multiple comparison were used (**C:** 3 x 10^-7^ M, 3 x 10^-6^M, 3 x 10^-5^M; **D:** 3 x 10^-7^, 3 x 10^-5^M; **E**, **F**: all concentrations). Sample sizes: n = 6 for 3 x 10^-7^M, n = 7 for 3 x 10^-6^M, n = 4 for 3 x 10^-5^M, n = 7 for 3 x 10^-4^M).

In the next set of experiments, we wanted to determine whether the effects of oxotremorine could be antagonized by the addition of scopolamine (Fig. 5A). We bath applied a dose of scopolamine (3 x 10^-7^M) that had minimal effects on the isolated CNS together with a concentration of oxotremorine (10^-5^M) that evoked significant effects on motor output (Fig. 3). The increase in event frequency normally observed in oxotremorine was blocked in the presence of scopolamine (Fig. 5A, B, n = 4). To confirm that these preparations were actually responsive to oxotremorine in the absence of scopolamine, oxotremorine was applied alone after washing out both drugs. Although the overall effects were not significant, all preparations showed increases in the frequency of fictive headsweeps and posterior bursts during this second oxotremorine application (Fig. 5C), confirming that scopolamine’s effects are at least partially reversible.

**Figure 5.**
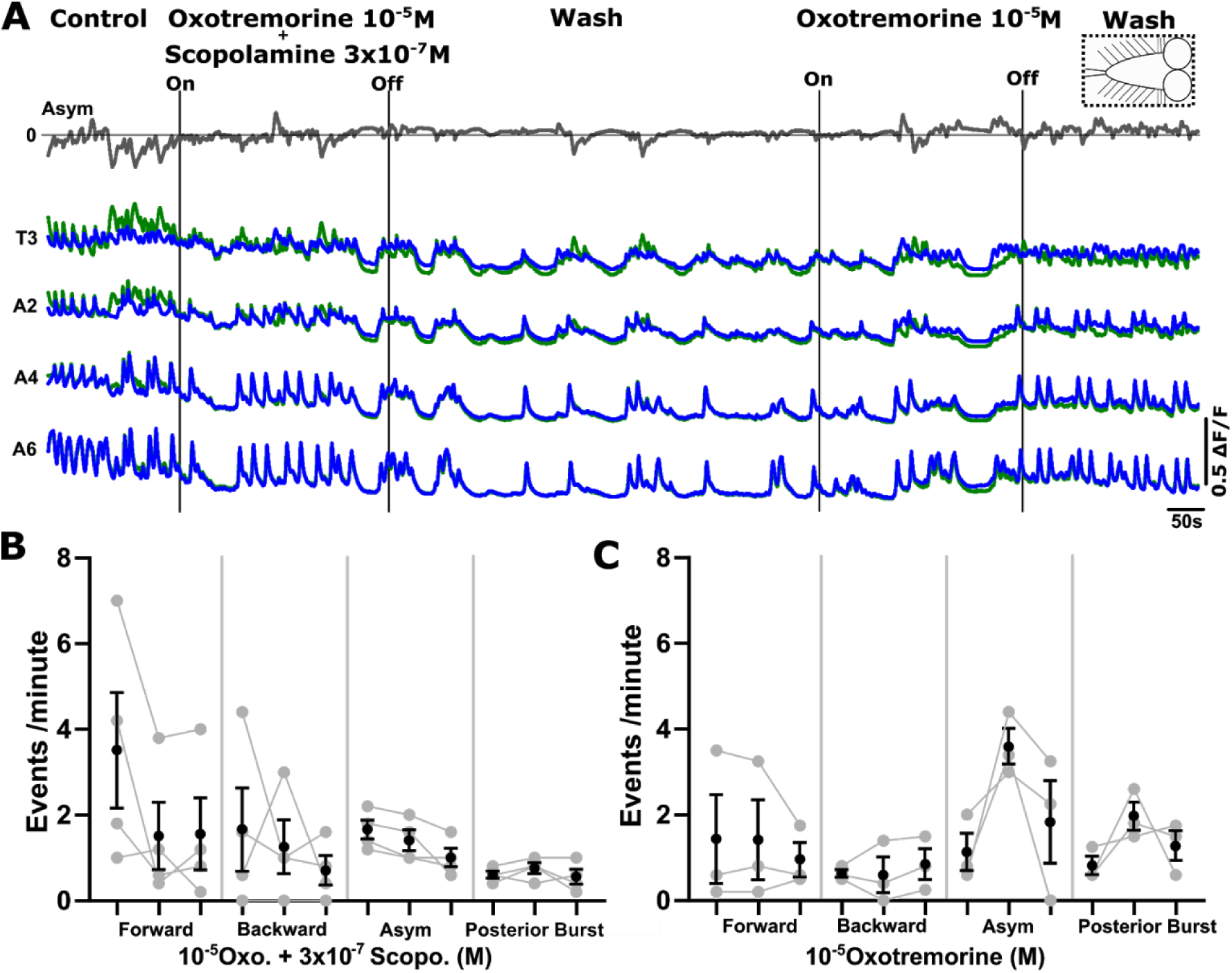
Low concentrations of scopolamine antagonize the effects of oxotremorine in the intact central nervous system. **A)** Ca^2+^ activity before during and after bath application of 3 x 10^-7^M scopolamine and 10^-5^M oxotremorine at same time (left), followed by wash then application of 10^-5^M oxotremorine alone (right). CNS is intact (dashed box at the top right). **B)** Pooled data showing no significant change in frequencies of fictive behaviours produced in intact CNS preparations during dual application of agonist and antagonist (n = 4, Friedman test with Dunn’s post hoc multiple comparison). **C)** Pooled data showing response to oxotremorine after wash of scopolamine; (n = 3, Friedman test with Dunn’s post hoc multiple comparison).

After confirming that manipulating mAChr signalling was an effective means to induce or suppress rhythm generation in this system generally, we then examined the effects of modulating mAChR signalling on isolated regions. Previous work has shown that the larval VNC is able to produce rhythmic motor patterns without brain lobes. To determine how signalling through mAChRs modulates activity generated intrinsically in the VNC, we surgically removed brain lobes, leaving the subesophageal zone (SEZ), thorax and an abdomen intact, then recorded activity before, during and after bath application of oxotremorine (Fig. 6A). We confirmed that motor neurons were indeed active in a subset of preparations using nerve root recordings and that hemisegmental Ca^2+^ signals were representative as in intact CNS preparations (data not shown). We then used Ca^2+^ imaging data for all subsequent analyses (Fig. 6A - D). When the brain lobes were removed, preparations became biased towards fictive forward waves with a subset of preparations being able to produce fictive backwards waves. Fictive headsweeps were not observed (compare Fig. 6A, top ‘Asym’ line to equivalent measure in Fig. 3A). Brain lobe ablated preparations showed a significant increase in fictive forward wave frequencies upon application of 10^-4^M and 10^-5^M oxotremorine (p < 0.05) (Fig. 6C). There was no significant effect on fictive backward wave frequencies at any concentration of oxotremorine (data not shown). In these preparations, we also observed a reversible increase in overall baseline fluorescence in 10^-4^M and 10^-5^M oxotremorine (p < 0.001, p < 0.05, respectively, Fig. 6D); baseline changes in fluorescence were not observed in oxotremorine in intact CNS preparations (Fig. 3A). In brain lobe excised preparations, the effects of 10^-5^M oxotremorine (Fig. 6) were antagonized with low concentrations of scopolamine (3 x 10^-7^M), as observed in intact CNS preparations (S1A, B).

**Figure 6.**
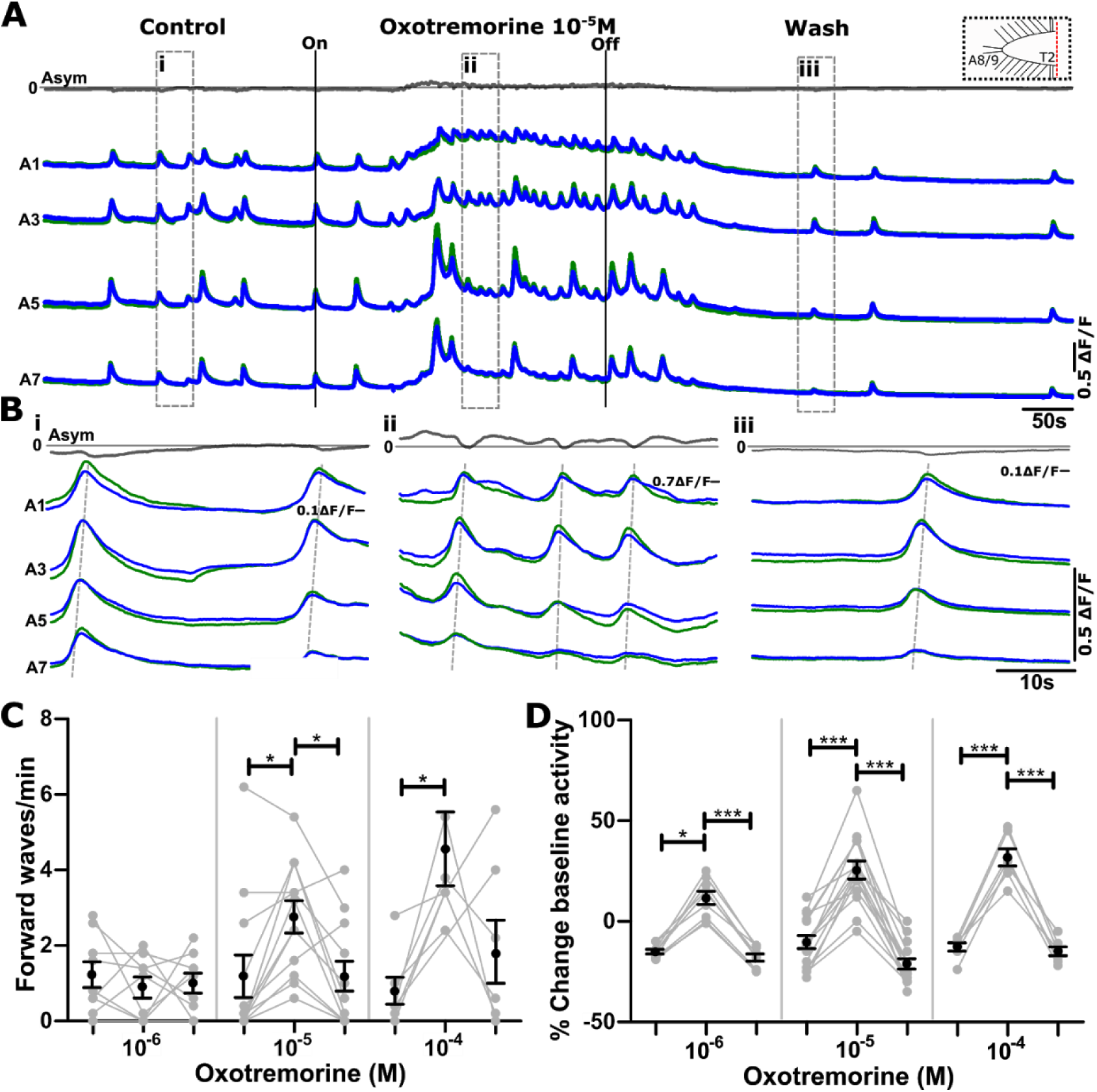
Oxotremorine increases frequency of forward waves and raises baseline Ca^2+^ levels in preparations without brain lobes. **A**) Ca^2+^ signals before, during and after application of 10^-5^M oxotremorine in a preparation without brain lobes (dashed box in upper right). **Bi)** Expanded view of Ca^2+^ signals in control period, **Bii)** expanded view of Ca^2+^ signals in oxotremorine, **Biii)** expanded view of Ca^2+^ signals in wash period. **C**) Mean +/− S.E.M fictive forward waves per minute produced before, during and after application of different concentrations (10^-6,-5,-4^M) of oxotremorine. Backward waves were sporadic in these preparations and no change in backward wave frequency was observed (data not shown). Bilateral asymmetries and posterior end bursts were not observed in these preparations in any conditions. **D)** Summary of baseline fluorescence changes before, during and after application of oxotremorine. Asterisks indicate significant differences amongst groups (*p < 0.05, **p < 0.005, ***p < 0.0001; Friedman test with Dunn’s post hoc multiple comparison tests were used). Sample sizes: **C, D**: n = 9 for 10^-6^M, n = 12 for 10^-5^M and n = 7 for 10^-4^M.

Next we set out to determine if anterior thoracic segments could produce fictive motor patterns without abdominal segments. In these experiments, we ablated the abdominal segments of the ventral nerve cord and recorded Ca^2+^ activity changes. In these preparations, the network was biased towards bilateral asymmetries (Fig. 7 A, B), no obvious wave-like activity was observed, but a subset of preparations did show periods of bilaterally synchronised bursts of activity (data not shown). The frequency of fictive headsweeps increased in a dose dependent manner in oxotremorine (10^-4^M: p < 0.01, 10^-5^M: p < 0.05, 10^-6^ M: non-significant, but trending towards increase). A non-significant trend towards an increase in baseline fluorescence was also observed at all concentrations of oxotremorine (Fig. 7D). This was followed by decreases in baseline fluorescence in the wash period (10^-4^M: p < 0.05, 10^-5^M: p < 0.05).

**Figure 7.**
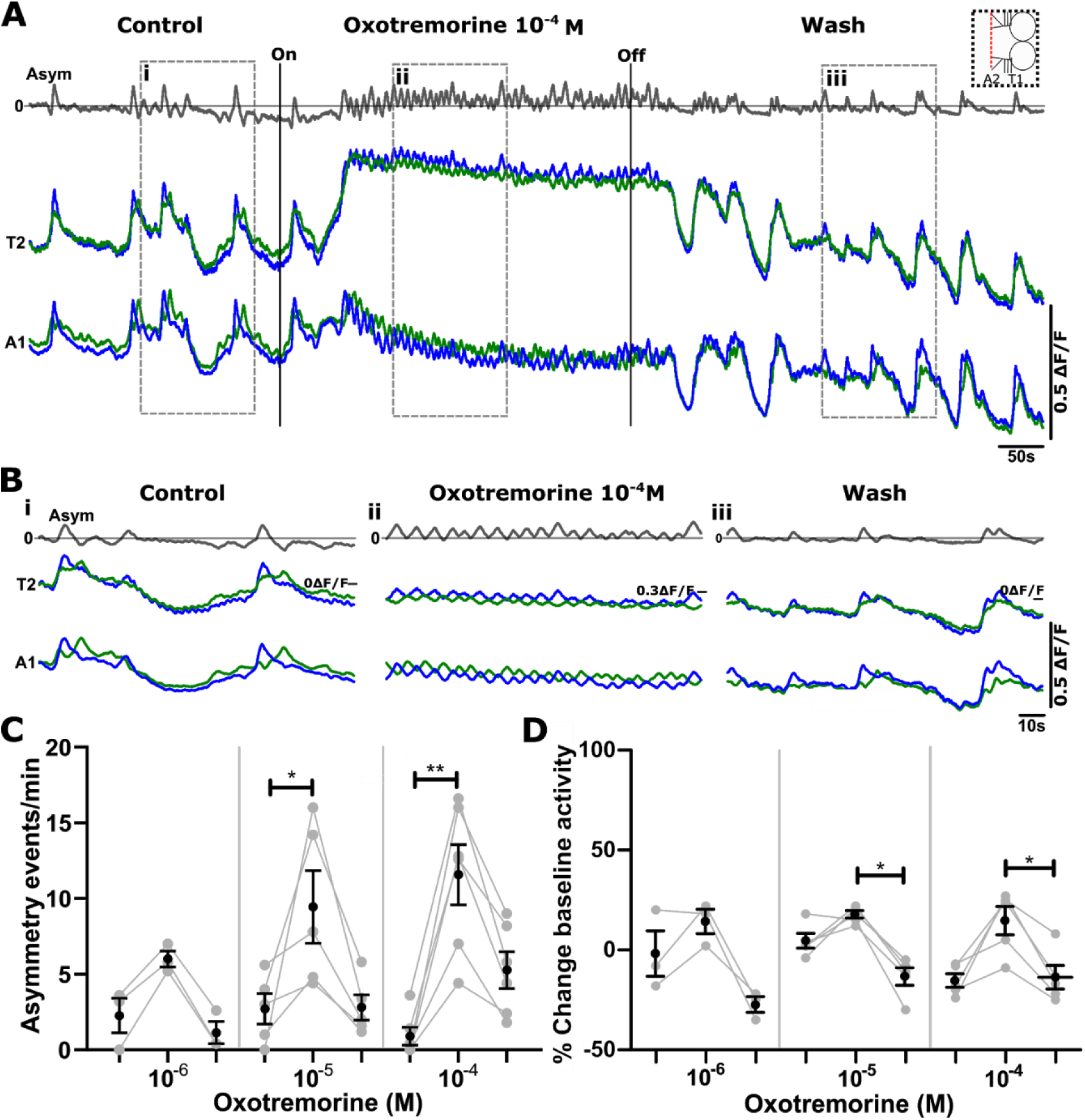
Oxotremorine increases the frequency of bilateral asymmetries and modulates baseline Ca^2+^ levels in preparations without abdominal segments. **A**) Ca^2+^ signals in thoracic and anterior abdominal segments before during and after oxotremorine in a preparation with posterior abdominal segments removed (schematic of preparation in upper right) **Bi**) Expanded view of Ca^2+^ signals in control period, **ii**) expanded view of Ca^2+^ signals during oxotremorine 10^-4^ M application, **iii**) expanded view of Ca^2+^ signals during wash period. **C**) Mean +/− S.E.M bilateral asymmetries per minute before, during and after application of three different concentrations of Oxotremorine (10^-6,-5,-4^M). Fictive forward and backward waves were not observed in any condition. **D)** Summary of baseline changes in Ca^2+^ fluorescence. Asterisks indicate significant differences amongst groups (*p < 0.05, **p < 0.005; Friedman test with Dunn’s post hoc multiple comparisons). Sample sizes: n = 3 for 10^-6^ M, n = 4 for 10^-5^M and n = 5 for 10^-4^M).

To further examine mAChR mediated effects on different parts of the VNC, we ablated brain lobes and all segments anterior to thoracic segment T2 while also transecting the posterior abdomen at abdominal segment A4 (Fig. 1Eiv). We were then able to simultaneously record activity from both anterior and posterior segments of the VNC in isolation from each other (Fig. 1Ev). Isolated anterior abdominal segments showed a reversible increase in baseline fluorescence (Fig. 8A, C, p < 0.01) but did not produce any rhythmic activity (Fig. 8A, B). In contrast, isolated posterior segments all showed reversible changes in baseline fluorescence as well as reversible increases in rhythmic activity in oxotremorine (Fig. 8D-F, p < 0.05).

**Figure 8.**
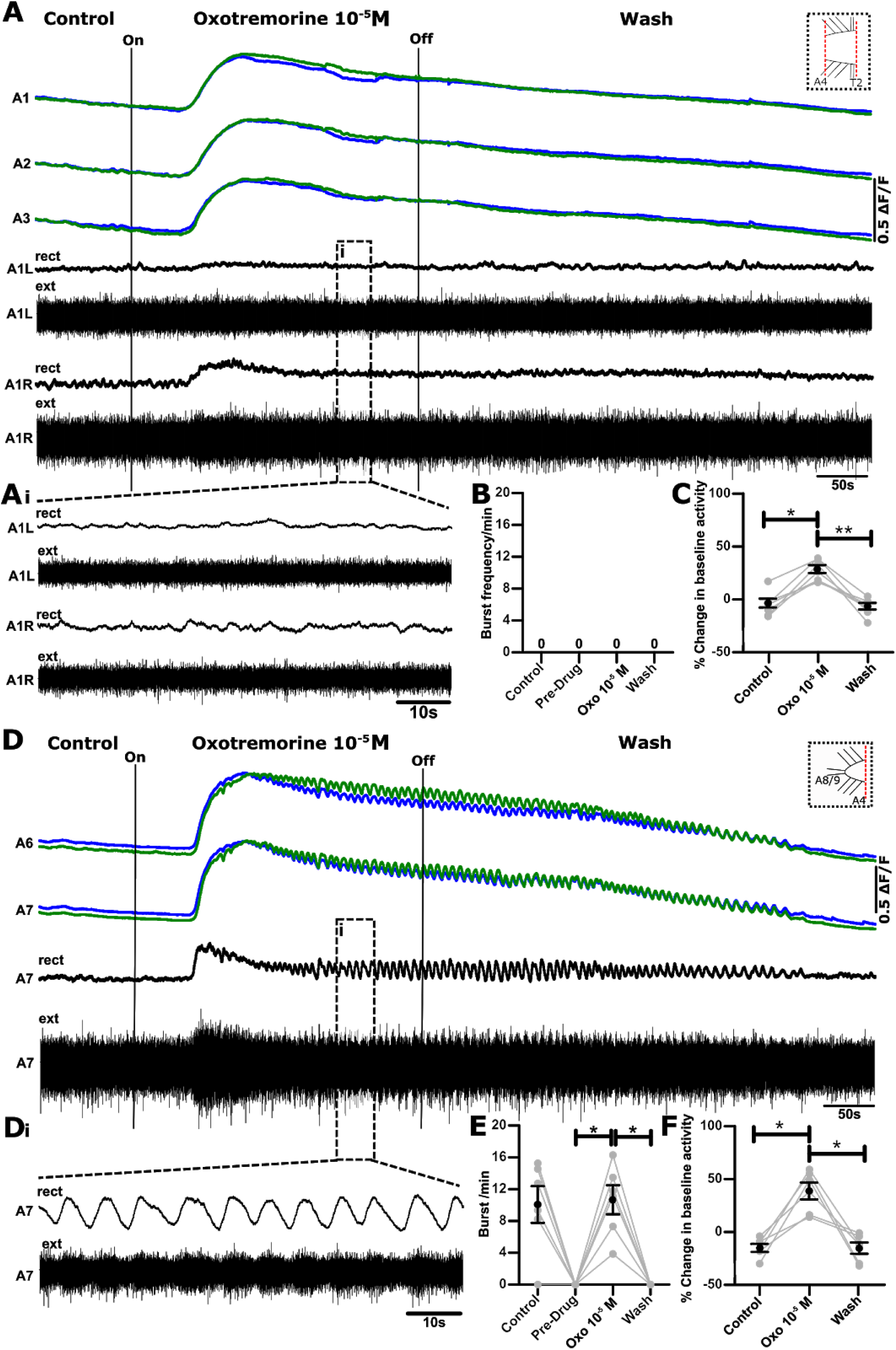
Oxotremorine triggers bursting in isolated posterior abdominal segments but not in isolated anterior abdominal segments. **A)** Example of simultaneous Ca^2+^ imaging (green and blue traces) and paired nerve root recordings (black traces) from isolated anterior abdominal segments before, during and after oxotremorine application (T2-A4, see schematic in upper right). Extracellular nerve root signals recorded from each electrode were rectified and smoothed by filtering with a moving average filter with a time constant of 0.9 s (rect). **Ai**) Expanded view of region in dashed boxes. **B, C)** Mean +/− S.E.M bursts per minute and baseline fluorescence in control, oxotremorine and wash, respectively. No bursts were detected in any condition (n = 7, see methods for burst criteria); however, all preparations showed reversible changes in baseline fluorescence. **D**) Ca^2+^ signals and single nerve root recording from posterior abdominal segments (A5-A8/9, see schematic in upper right). **Di)** Expanded view of black dashed boxes in **D.** Note rhythmic activity is not present in Ai. **E, F)** Mean +/− S.E.M bursts per minute and baseline fluorescence in control, oxotremorine and wash, respectively (n = 7). Asterisks indicate significant differences amongst groups (*p < 0.05, **p < 0.005; Friedman test with Dunn’s post hoc multiple comparisons).

Next, we aimed to investigate if scopolamine could inhibit motor pattern generation in transected parts of the VNC (Fig. 1Eii, iii, v). In three types of transected preparations (brain lobes removed, posterior abdomen removed, all anterior segments removed), application of low doses of scopolamine (3 x 10^-6^ M) inhibited or abolished rhythmic activity (p > 0.05, p < 0.005, p < 0.05, respectively, Fig. 9). All preparations showed at least partial recovery of rhythmicity in wash periods, confirming that rhythm generating circuits retained functionality after scopolamine application.

**Figure 9.**
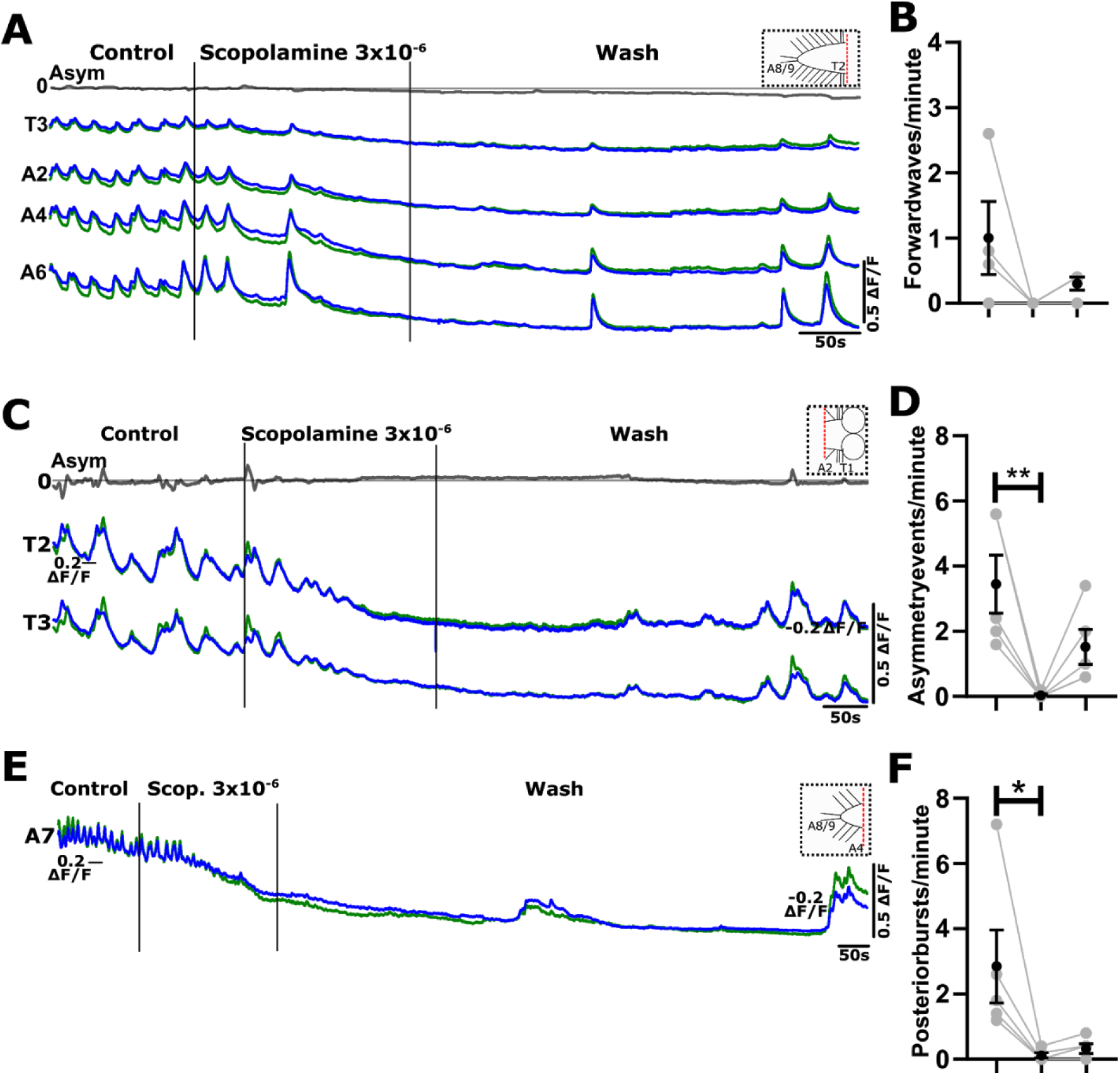
Scopolamine inhibits bursting in different ablated regions of the central nervous system. **A-F)** Examples of Ca^2+^ signals from three different ablated parts of CNS (top right corners of **A**, **C** and **E** depict type of preparation). **B, D, F)** Mean +/− S.E.M events per minute before during and after scopolamine. Note complete suppression of rhythmic activity and partial recovery in wash. Sample sizes (B, D, F): n = 4, 5, 5 respectively). Asterisks indicate significant differences amongst groups (*p < 0.05, **p < 0.005; Friedman test with Dunn’s post hoc multiple comparisons).

Finally, to determine whether midline connections are required to produce rhythmic activity, we ablated one entire side of the CNS (Fig. 1Evi) and measured responses to oxotremorine (Fig. 10A, B). Bath application of oxotremorine to these preparations triggered an increase in baseline fluorescence and largely synchronous rhythmic bursting in anterior regions, but did not induce fictive forward or backward waves. (Fig. 10A, n = 3).

**Figure 10.**
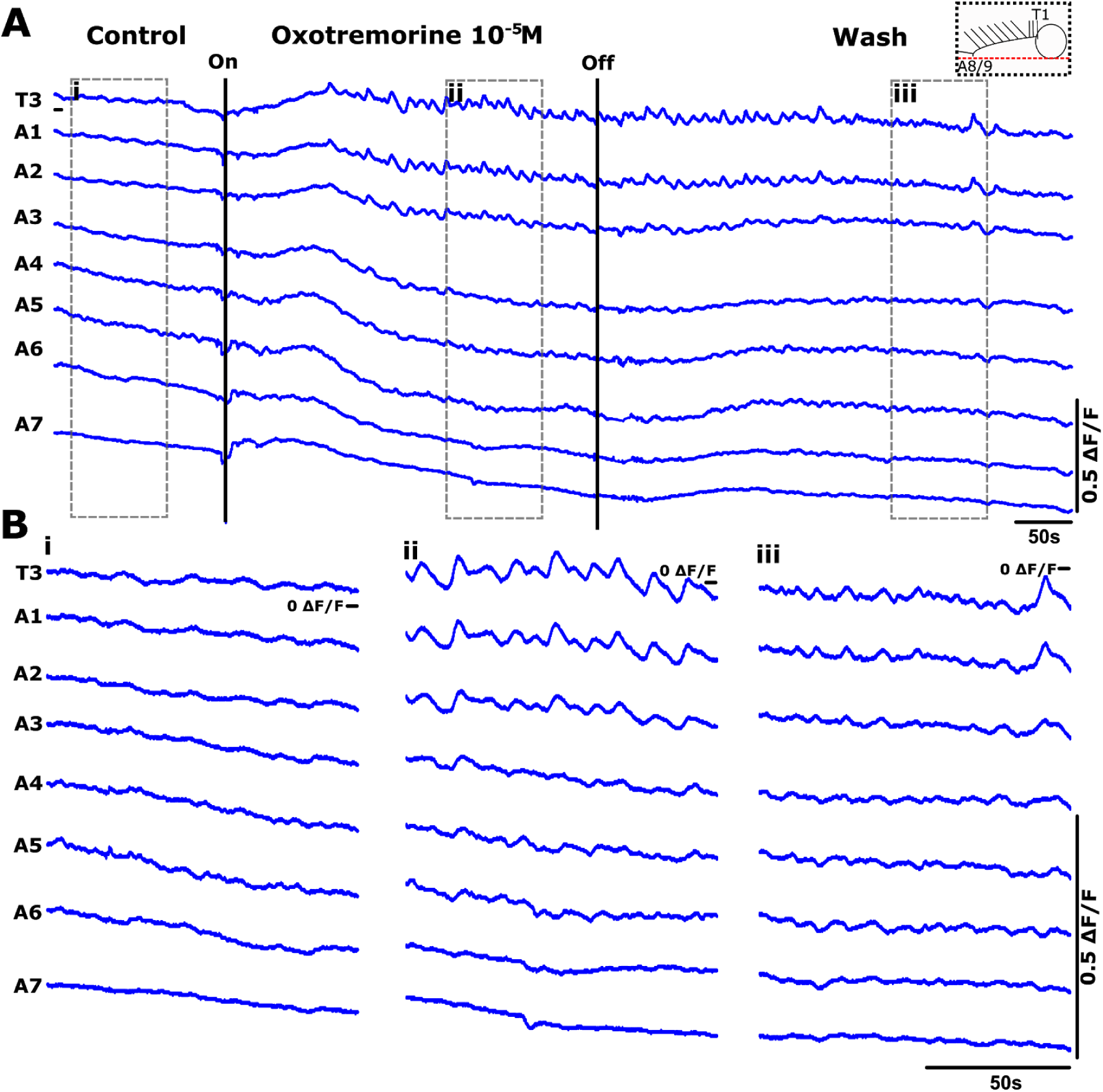
Oxotremorine triggers bursting in anterior regions in longitudinally hemi-sected preparations. **A)** Representative traces of Ca^2+^ signals from a longitudinally hemi-sected preparation before during and after oxotremorine application (schematic of the preparation shown in top right). **B)** Expanded view of Ca^2+^ signals in (i) control period (ii) 10^-5^ M oxotremorine and (**iii**) wash period. Note bursts of activity in anterior regions. Results representative of 3 preparations.

A schematic showing a summary of results is shown in Figure 11. Overall, our results suggest the presence of multiple gradients of rhythmogenicity, with separate rhythm generating modules present in anterior brain and thoracic regions as well as in posterior abdominal segments. In contrast, anterior abdominal regions appear to have little or no mAChR dependent rhythmogenic properties. Midline connections are not required for rhythm generation in anterior regions, but may play a critical role for rhythm generation in posterior segments.

**Figure 11.**
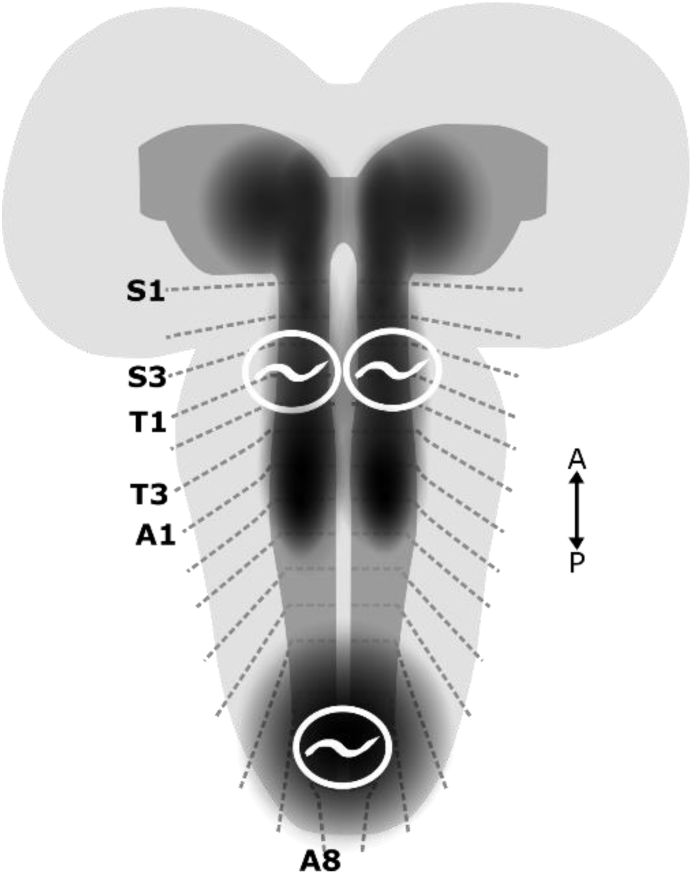
Proposed location of mAChR dependent rhythm generating regions in the *Drosophila* larval CNS. Schematic diagram showing larval CNS. Dashed grey lines point to approximate locations of neuromeres in SEZ, thorax and abdomen; neuropil regions are shaded in dark grey. Regions showing evidence of mAChR dependent rhythmogenicity are shaded in black with white oscillator symbol. Note that while anterior abdominal regions are not strongly rhythmogenic on their own, they can initiate backward waves, suggesting some potential for rhythm generation.

## Discussion

Here we used a combination of electrophysiology, Ca^2+^ imaging and pharmacology in isolated CNS preparations to explore the architecture of rhythm generation in the *Drosophila* larval locomotor system. We pharmacologically manipulated mAChR signalling and measured the resulting effects on a variety of fictive behaviours underlying larval locomotion, including forward and backward waves and turning/headsweeping behaviour. The mAChR agonist oxotremorine, did not strongly affect the frequency of forward or backward fictive locomotion, but did potentiate fictive headsweeps and bursting in posterior abdominal regions. The mAChR antagonist scopolamine blocked the effects of oxotremorine and inhibited rhythmic activity when applied alone. In reduced preparations, oxotremorine potentiated rhythm generation in isolated distal regions of the VNC while scopolamine abolished rhythmic activity in the same types of preparations. Overall, these results suggest that mAChR signalling is required for generating larval locomotor rhythms and provides initial evidence for the presence of multiple independent mAChR-dependent rhythm generating modules in distal regions of the larval CNS.

To manipulate muscarinic signalling, we used a pharmacological approach in which we bath applied different concentrations of the mAChR agonist, oxotremorine (10^-4^ to 10^-6^ M) and mAChR antagonist, scopolamine (3 x 10^-4^ to 3 x 10^-7^ M). We chose these concentrations because they have been shown to activate *Drosophila* mAChRs (and not nAChRs) in heterologous expression systems (Collin et al., 2013; Ren et al., 2015) and because they are equivalent or lower to those used in other studies of vertebrate (Miles et al., 2007; Gross and Bloomquist, 2018) and arthropod locomotor networks (Ryckebusch and Laurent, 1993; BÜSchges et al., 1995; Johnston and Levine, 1996; Cattaert and Birman, 2001; Buhl et al., 2008; Rillich et al., 2013). Similarly, the concentrations of antagonists used are known to block *Drosophila* mAChRs and at the lowest concentration, are primarily selective for A- and C-type receptors while not strongly affecting B-type receptors (Collin et al., 2013; Ren et al., 2015). Critically, the effects of oxotremorine could be fully antagonized by co-application of low (i.e. < 10^-6^ M) doses of scopolamine, suggesting that oxotremorine is indeed acting through mAChR receptors and not through off-target mechanisms.

Taking a pharmacological approach in an *in vitro* preparation limited our ability to precisely target specific receptor sub-types (drugs specifically targeting *Drosophila* subtypes are not available). While it is possible to use RNA interference (RNAi) to suppress expression of specific mAChR subtypes (Silva et al., 2015; Malloy et al., 2019), these genetic approaches manipulate mAChR expression over developmental time scales. This raises the possibility that any phenotypes observed in RNAi studies are the end result of developmental compensation in response to mis-expression of receptors. A major advantage of our pharmacological approach is that it allowed us to acutely manipulate mAChR signaling on the time scale of minutes, thereby providing a clear view of receptor function without the confounding variable of developmental compensation.

Previous work in *Drosophila* larvae has shown that longer-term pharmacological and genetic manipulations of mAChR activity can lead to non-intuitive changes in larval behaviours. Feeding animals muscarine for 24 hours resulted in lower forward step frequency, whereas feeding them antagonists had a similar effect (Malloy et al., 2019). In another study, larvae fed mAChR antagonist atropine showed relatively normal locomotion but failed to navigate away from noxious odors (Silva et al., 2015). Interestingly, genetic knockdown of specific subtypes of mAChR using RNAi gives a different set of results. Suppressing expression of A- and C-type receptors in a variety of cell populations actually leads to an increase in forward step frequency (Malloy et al., 2019). One explanation for the mismatch in behaviour effects between pharmacological and genetic manipulation observed in (Malloy et al., 2019) could be that both manipulations are occurring over relatively long periods (1 - 4 days). On this time scale, compensatory homeostatic mechanisms could be engaged (or over engaged) in response to the manipulations, thereby complicating interpretation of resulting behaviours.

In previous work, acute application of oxotremorine and other muscarinic agonists triggered peristaltic waves of activity in semi-intact larval preparations and potentiated sensori-motor circuitry; conversely, mAChR antagonists generally suppressed activity (Cattaert and Birman, 2001; Malloy et al., 2019). In these studies, motor activity was measured by recording excitatory junctional potentials (EJPs) at the neuromuscular junction and sensory feedback was intact in semi-intact preparations. Recent work has also examined activity in single nerve roots in isolated CNS preparations in response to mAChR agonists and antagonists (Gross and Bloomquist, 2018). Overall, this previous body of work largely agrees with our findings, namely that muscarinic agonists potentiate locomotor circuit activity and antagonists suppress activity. Although insightful, these previous electrophysiological studies could only provide limited views of the overall activity of the larval locomotor network because they only recorded activity in 1 or 2 body segments at a time.

Our imaging experiments allowed us to measure fictive locomotion across most VNC segments through a combination of Ca^2+^ imaging and simultaneous extracellular nerve root recordings. First, we wanted to test how bath application of the agonist oxotremorine would affect spontaneous rhythmic activities produced by the isolated larval CNS. The recordings were at first difficult to interpret, as there were no strong effects on overall forward or backward wave frequency. Further analysis revealed that the main effect of oxotremorine was actually to potentiate bilateral asymmetries (fictive head sweeps) in thoracic regions and posterior bursting in posterior abdominal regions. Bath application of scopolamine reliably suppressed multiple fictive motor programmes, with especially strong effects on fictive head sweeps. These results suggest that mAChR signalling is critically important for generating rhythmic motor patterns in the larval VNC that require intersegmental and/or bilateral coordination. These results are similar to those obtained in the locust flight CPG (Buhl et al., 2008), various insect walking CPGs (Ryckebusch and Laurent, 1993; BÜSchges et al., 1995; Johnston and Levine, 2002; Buhl et al., 2008) (Johnston and Levine, 1996). In vertebrate spinal cord, signalling through NMDA receptors appears to play a key role in raising cytosolic Ca^2+^ which then contributes to initiation of rhythmic activity in both intact and hemi-sected preparations (Guertin and Hounsgaard, 1998; Miles and Sillar, 2011; Hägglund et al., 2013). Signalling through mAChRs may be playing an analogous role in promoting rhythmogenesis in insect locomotor CPGs, with activation of A- and/or C-type mAChRs leading to release of intracellular Ca^2+^ stores through Phospholipase C triggered transduction pathways (Collin et al., 2013).

We noted in our initial experiments that the initiation of vigorous bouts of fictive head sweeps and posterior bursting appeared to suppress a preparation’s ability to generate wave-like activity. We reasoned that oxotremorine may be triggering two competing independent rhythm generators at either end of the network. To test this idea, we performed a series of ablation experiments aimed at localizing muscarinic dependent rhythm generating modules. In brain lobe ablated preparations, bilateral asymmetries were abolished, and forward wave frequency was increased. In these preparations, oxotremorine triggered posterior bursting which, in turn, triggered forward waves on nearly every cycle. These effects were completely blocked by the addition of scopolamine and recovered in the same preparations after wash period (S1). This suggests that a local muscarinic dependent rhythm generating mechanism exists in abdominal regions and this mechanism primarily triggers forward waves. These results also provide indirect evidence that fictive head sweep circuitry suppresses the generation of forward waves (in agonist, in the absence of fictive headsweeps, forward waves dominate). In the next set of experiments, we ablated abdominal regions and measured activity in preparations with just brain lobes and thoracic regions remaining. Application of oxotremorine triggered dose dependent increases in the frequency of bilateral asymmetries; however, even in the absence of agonist, preparations generated bilaterally asymmetric activity. These results suggest that a separate rhythm generating module in anterior regions underlies fictive head sweeps. Interestingly, recent work has independently suggested that continual lateral oscillations of anterior regions during navigation are driven by neuronal oscillators which could be subject to neuromodulation (Wystrach et al., 2016). The results presented here corroborate this idea and provide an entry point to identifying the circuit components underlying headsweep oscillations.

We continued systematically removing parts of the VNC to determine which areas are rhythmogenic and which are not. To confirm the presence or absence of any possible type of rhythmic activity, wherever possible, we performed both panneuronal imaging and extracellular nerve root recordings simultaneously. One striking result was that middle sections of the VNC (A1 - A5) did not produce any rhythmic activity even in oxotremorine (Fig. 8A). They did, however, respond to oxotremorine with a raised baseline Ca^2+^ signal that reversed in wash conditions, demonstrating that the segments were viable. Conversely, isolated posterior segments from the same preparations were highly rhythmogenic and showed regular bursting in oxotremorine (Fig. 8D). These results suggest that segments in the VNC are not all equally capable of generating rhythmic activity when isolated. This is not consistent with existing models of the larval locomotor system that assume that each hemi-segment contains an identical oscillator that is then coupled to other oscillators in every adjacent segment (Gjorgjieva et al., 2013). Our work suggests that the larval VNC may be organized somewhat differently, with strongly rhythmogenic oscillators at distal ends separated by segmentally organized circuitry that primarily promotes wave propagation. Our results do not definitively rule out the presence of oscillators in medial hemi-segments. Oscillators may be present but simply less rhythmogenic. They could also be present but triggered by different modulatory mechanisms. indeed, the fact that preparations without thoracic regions and brain lobes can initiate bouts of backward waves suggests the presence of ‘latent’ oscillators in anterior abdominal regions (Pulver et al., 2015). This is partially in line with previous work in vertebrates showing the presence of rostral-caudal gradients of rhythmogenicity in spinal circuitry (Kjaerulff and Kiehn, 1996; Wiggin et al., 2012, 2014; Hägglund et al., 2013); however, in vertebrates, the gradient is unimodal with rhythmogenicity steadily declining rostral to caudal. In contrast, in the soft-bodied leech, all segments appear equally capable of producing crawling motor programmes in isolation in the presence of dopamine, with both descending neurons and local circuits providing substrates for inter-segmental coordination (Puhl and Mesce, 2008).

In the intact CNS, the muscarinic antagonist scopolamine reduced the frequency of fictive motor patterns but did not completely abolish rhythmic activity (Fig. 4). In contrast, relatively low doses of scopolamine completely abolished endogenous rhythmic activity in brain ablated isolated anterior and isolated posterior segments (Fig. 9). These data suggest that muscarinic signalling is indeed required for rhythm generation in distal rhythm generators. This complements previous work showing that pharmacological block of nicotinic acetylcholine receptors (nAChR) abolishes all rhythmic synaptic activity (Rohrbough and Broadie, 2002) and calcium oscillations (J. Booth, Pulver laboratory, personal communication), suggesting that fast excitation generally, and specifically signalling through nAChRs is also required for rhythm generation. Conversely, bath application of picrotoxin leads to seizure-like activity followed by eventual collapse of rhythmic activity (Streit et al., 2016) suggesting that inhibition generally, and specifically inhibition mediated by GABAA and/or Glutamate-gated chloride channels, is also required for rhythm generation. Excitatory and inhibitory interneurons with critical roles in wave pattern generation have recently been identified in the larval CNS (Iyengar et al., 2011; Fushiki et al., 2016; Yoshikawa et al., 2016; Clark et al., 2018; Kohsaka et al., 2019; Zarin et al., 2019); however, further signalling requirements for rhythm generation in this system remain unclear. In particular, the contribution of rhythmogenic ionic currents (Harris-Warrick, 2002; Gao et al., 2018) is largely unexplored and could be the focus of future work.

In a final set of experiments, we asked whether bilaterally asymmetric motor patterns in thoracic regions require connections across the midline. We longitudinally transected the entire CNS and recorded endogenous and oxotremorine evoked activity. These preparations were not able to produce wave-like activity or posterior bursting; however, they were able to produce synchronous activity in thoracic segments in the presence of oxotremorine (Fig. 10). These results suggest that bilateral asymmetric activity arises from interactions between independent rhythm generating modules located on either side of the CNS, as is the case in vertebrate spinal cord (Hägglund et al., 2013). These results also tentatively suggest that contralateral connections may be required for rhythm generation in posterior segments. Importantly, the lack of rhythmic activity in posterior regions in hemi-sected preparations should not be regarded as conclusive evidence for critical contralateral connections, especially given the difficulty of making precise longitudinal cuts at the narrow posterior tip of the VNC.

Overall, this work identifies mAChRs as key components of rhythm generation in the *Drosophila* larval CNS. We provide evidence for presence of mAChR dependent rhythm generating modules in distal regions of the larval VNC. These rhythmogenic regions are separated by medial segments with limited rhythmogenic ability (Fig. 11). Previous work suggests that mAChRs are widely distributed in the brain and VNC of larvae (Silva et al., 2015); however, it is unclear exactly how mAChRs are distributed amongst identified interneurons. Future work aimed at carefully characterizing the spatial distribution of mAChRs amongst identified cell types in the larval VNC could be a productive next step towards identifying the interneuron cell types at the core of rhythm generating circuits in the larval VNC.

## Acknowledgments

We would like to thank Dr. Maarten Zwart (University of St Andrews) and Dr. Andrew Dacks (West Virginia University) for constructive comments on an earlier version of this manuscript. We also thank Karen Hibbard (Janelia Research Campus) for advice on fly lines.

---

**Figure supplement 1.**
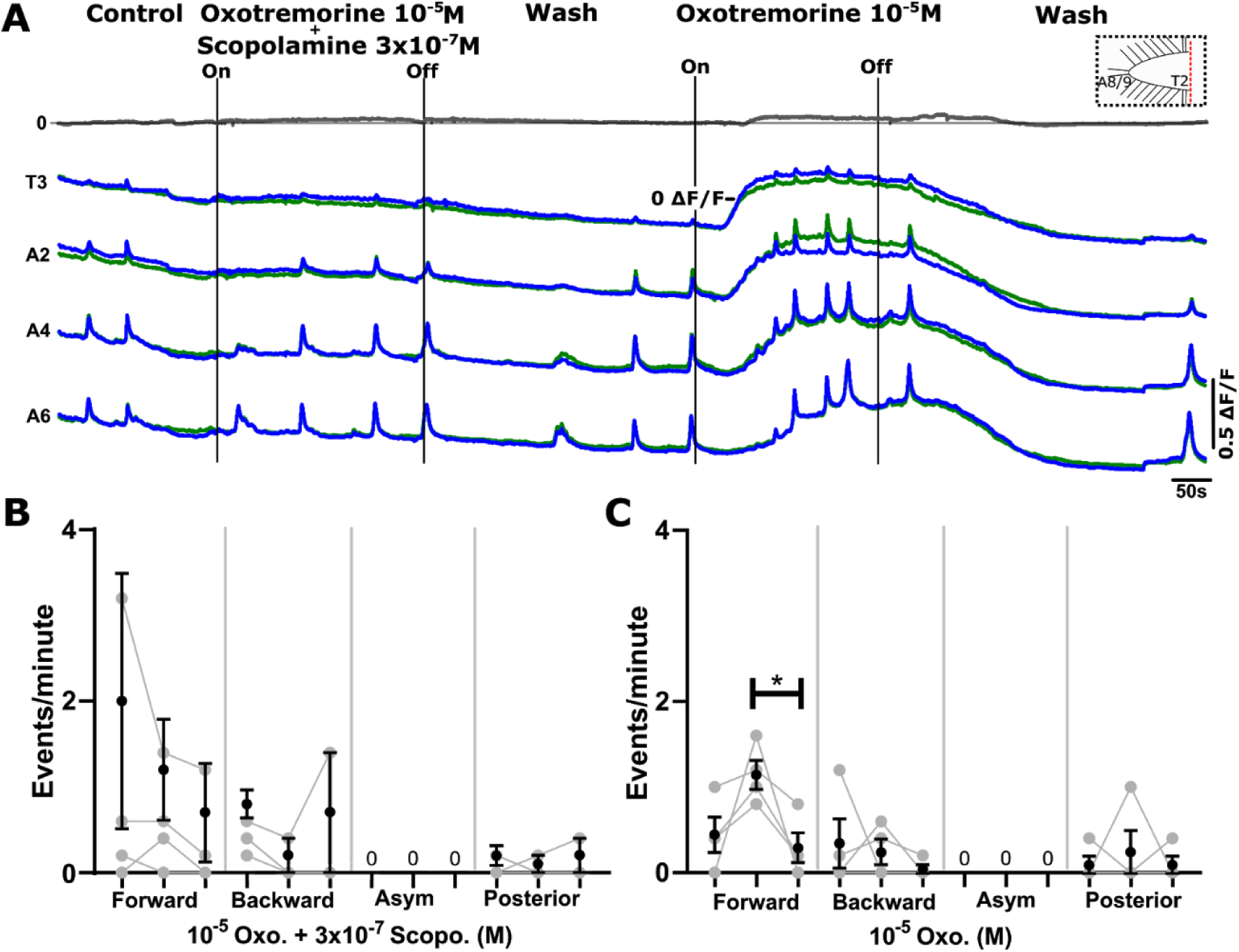
Low concentrations of scopolamine antagonize effects of oxotremorine in preparations without brain lobes. **A**) Representative examples of Ca^2+^ signals during co-application of 10^-5^M oxotremorine and 3 x 10^-7^M scopolamine (left section); following a wash period (middle section), 10^-5^M oxotremorine was applied alone (right section). Note lack of change in frequency and baseline during co-application and increase in frequency and baseline fluorescence in oxotremorine alone. **B**) Mean +/− S.E.M events per minute before, during and after co-application for 4 motor programmes. **C)** Same as in B but in oxotremorine alone after wash period. Samples sizes B, C: n = 4. Asterisks indicate significant differences amongst groups (*p < 0.05; 1 way ANOVA, Bonferroni post-hoc test).

